# Linear and Neural Network Models for Predicting N-glycosylation in Chinese Hamster Ovary Cells Based on B4GALT Levels

**DOI:** 10.1101/2023.04.13.536762

**Authors:** Pedro Seber, Richard D. Braatz

## Abstract

Glycosylation is an essential modification to proteins that has positive effects, such as improving the half-life of antibodies, and negative effects, such as promoting cancers. Despite the importance of glycosylation, predictive models have been lacking. This article constructs linear and neural network models for the prediction of the distribution of glycans on N-glycosylation sites. The models are trained on data containing normalized B4GALT levels in Chinese Hamster Ovary cells. The ANN models achieve a median prediction error of 1.39%, which is 10-fold smaller than for previously published models, and a narrow error distribution. We also discuss issues with other models reported in the literature. We provide all of the software used in this work, allowing other researchers to reproduce the work and reuse or improve the code in future endeavors.

## 1 Introduction

Glycosylation is a form of co-translational and post-translational modification that involves adding a glycan or glycans to proteins. When a glycan is added to the nitrogen of an asparagine or arginine, this process is called N-linked glycosylation. Glycosylation has many important functional and structural roles [1] [2] [3]. Improper glycosylation or deglycosylation, on the other hand, is associated with multiple diseases such as cancers [4], infections [5], and congenital disorders [6]. Many enzymes participate in the glycosylation process, and B4GALT1–B4GALT4 have been shown to be key contributors in multiple independent studies [7].

Due to its potential for diagnoses and treatments, glycosylation has been of significant interest to the biomedical and pharmaceutical industry, physicians, and patients. For example, increases in fucosylation, branching, and sialyation occur in many types of carcinoma [8]. Disialoganglioside is expressed by almost all neuroblastomas, and anti-disialoganglioside monoclonal antibodies have been successful against high-risk neuroblastoma in Phase I-III studies [9] [10]. Another example is poly-*α*2,8-sialyation, which increases the half-lives of antibodies without introducing tolerance problems [11]. On the other hand, the presence of glycans not produced by humans can be detrimental to a therapeutic. N-glycolylneuraminic acid, which is in some CHO-cell-derived glycoproteins [12], is immunogenic to humans [13].

Despite the importance of glycosylation for biotherapeutics and the many advances made in the field, such as the genetic engineering of CHO cells to increase the sialyation of glycoproteins [14], some challenges remain. Proteins have multiple glycosylation sites, and analyses need to take into account not only the glycan compositions but also where these glycans are located [8]. This structural diversity makes it difficult to explore specific functions of glycosylation [3]. Clinical laboratories struggle with analyzing patient glycosylation samples due to the complex equipment needed, which has limited the progress of personalized medicine [8].

Many computational models have been developed to assist researchers in better understanding and predicting glycosylation patterns. Some of the models, such as YinOYang [15] or O-GlcNAcPRED-II [16], are classifiers. In the context of glycosylation, these models may predict whether an amino acid is N-GlcNAcylated, or whether it is O-GlcNAcylated, for example. Other models, such as by Ref. [17], perform regression, that is, the models attempt to predict numerical values. In the context of glycosylation, a regression model may predict the numerical glycan distribution at a given glycosylation site.

In this work, we construct linear models and artificial neural network (ANN) regression models to predict glycan distribution in different glycosylation sites from normalized B4GALT levels in Chinese Hamster Ovary cells. The model construction procedures employ nested cross-validation and rigorous unbiased prediction error estimation. Our ANN models have 10-fold lower median prediction error than previously reported models. Open-source software is provided so that other researchers can reproduce the work or retrain the models as additional glycosylation data become available.

## 2 Materials and Methods

This section describes the datasets and methods for constructing the data-driven models. Details on how to run the software to make glycan distribution predictions or recreate the results in this article are provided in Supplementary Information.

### 2.1 Datasets

The models in this article are constructed using two experimental datasets.^1^ The larger dataset, from the Supplemental Data of Ref. [7], comprises distributions of glycans in nine different glycosylation sites in response to changes in the enzymes B4GALT1–B4GALT4 due to five types of knockouts. A smaller dataset, from Figures 3–5 and S2 of Ref. [18], comprises distributions of glycans over time in response to changes in the concentrations of nucleotide sugar donors. 20% of each dataset was separated for testing, with the remaining 80% used for cross-validation. The data from Ref. [7] are split into five groups according to the knockouts that are performed and the data from Ref. [18] into five groups according to the feed conditions.^2^ During each cross-validation fold and testing, the validation/testing groups are selected such that an entire knockout group was held out, and the other knockout groups are used for training. This procedure avoids test set leakage by ensuring the test set is sufficiently different from the training set.

### 2.2 Linear data-driven models

Linear data-driven models are constructed using elastic net, ridge regression, and partial least squares implemented in the Smart Process Analytics (SPA) software [19].^3^ For each data target, the best hyperparameters for the linear models are selected through cross-validation on four of these groups. The model and hyperparameters with the lowest cross-validation error are selected and the last group used to test that model’s performance.

### 2.3 Artificial Neural Networks (ANNs)

Artificial neural network models (specifically, multilayer perceptrons) are constructed for both datasets using PyTorch [20] and other Python packages [21] [22] [23]. We constructed models for 7 different layer configurations, 6 different learning rates (5e-2, 1e-2, 5e-3, 1e-3, 5e-4, 1e-4), and 4 different activation functions (ReLU, SeLU, tanh, tanhshrink), and plateau or cosine scheduling [24].^4^ The best hyperparameters for each glycan are determined by a grid search. The combination with the lowest cross-validation average loss is selected and, for each glycan, its performance is reported for an independent test dataset.

## 3 Results

### 3.1 Properly trained linear and ANN models display good performance on these datasets

All of the data-driven models constructed in this study for prediction of N-glycosylation are trained with hyperparameters selected by cross-validation. This section compares our models with the models in Ref. [25] (denoted by ANN-2 and ANN-3), which trained ANNs using the same data but selected the hyperparameters based on test set performance, a procedure that leads to overly optimistic model errors and issues with generalization [26] [27]. Thus, it should be noted that the real relative errors for the models from Ref. [25] are higher than what is reported in this work or Ref. [25]. Moreover, the training procedure of Ref. [25] had some restrictions, such as on the learning rate and activation function, that reduced the potential prediction accuracy of its models.

The predictions of the data-driven models are compared with experimental data in the test sets in Fig. 1, and the corresponding percent relative errors (PRE) are summarized in Table 1. The median prediction error for our ANN is 10-fold lower than for the previously published models. The mean and median prediction errors for ANN-2 and ANN-3 are higher than for our ANN, in spite of being selected based on their performance on the test set instead of on a validation set, indicating issues with the ANN training in Ref. [25].

**Figure 1:**
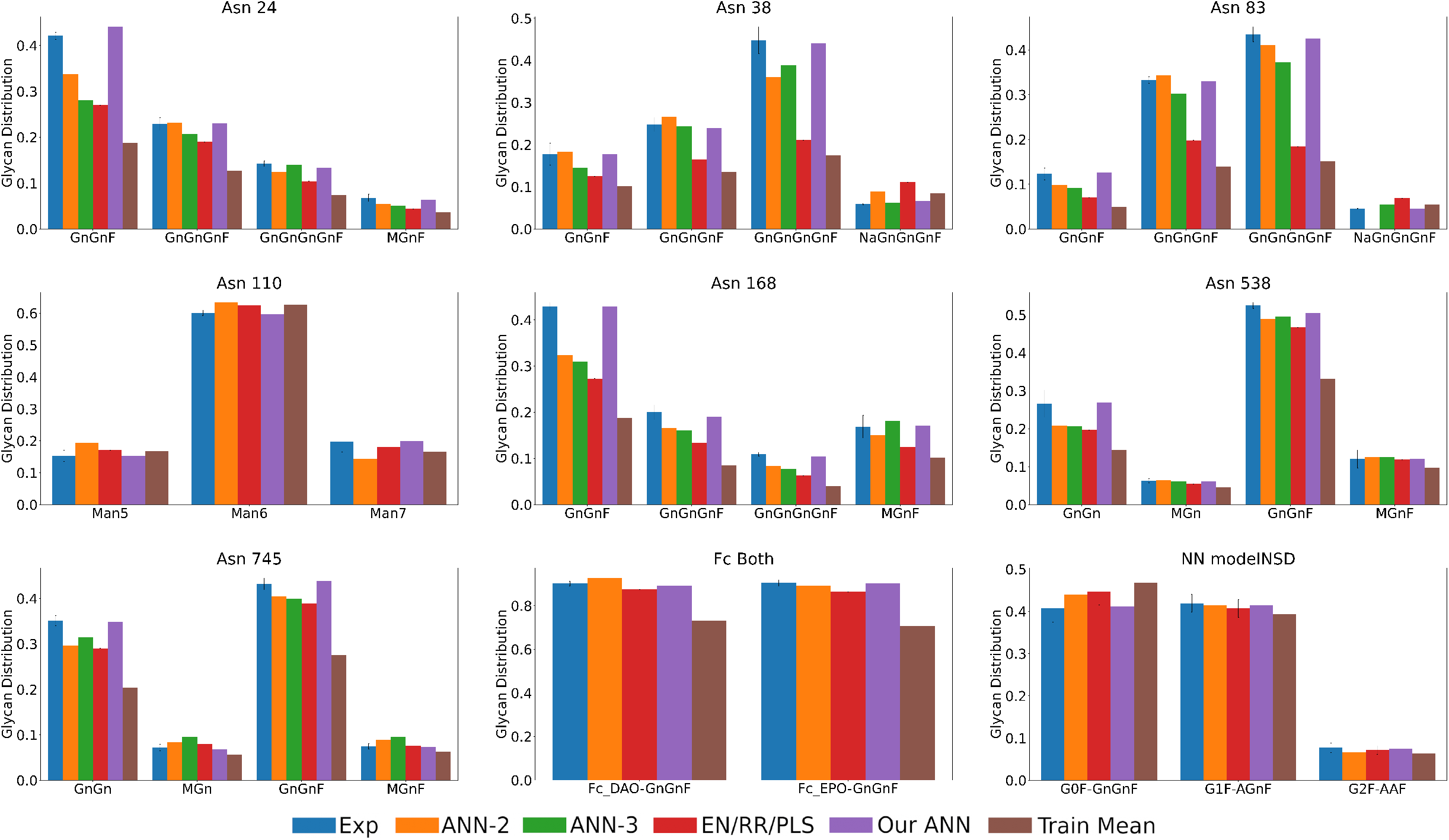
Glycan distributions predicted by different models from test data. Experimental results (“Exp”) came from Refs. [7] for the first 8 subplots and [18] for the last subplot (see Section 2.1). Models “ANN-2” and “ANN-3” came from Ref. [25]. Models “EN/RR/PLS” and “Our ANN” come from this work. “Train mean” is the mean of the training data (Refs. [7] and [18]). For each subplot, the bar orders and colors are the same, and each x-axis position represents a different glycan.

**Table 1:**
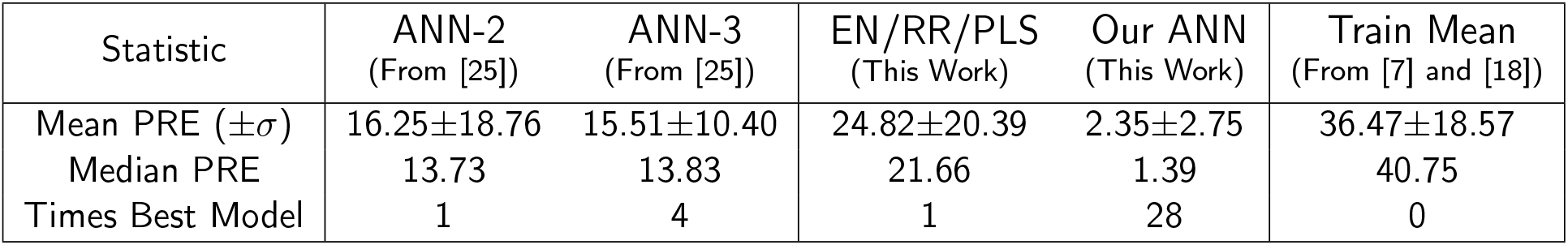
Mean and median percent relative errors (PRE) for different N-glycosylation models. Models “ANN-2” and “ANN-3” are from Ref. [25]. Models “EN/RR/PLS” and “Our ANN” come from this work. “Train mean” is the mean of the training data (Refs. [7] and [18]).

| Statistic | ANN-2<br>(From [25]) | ANN-3<br>(From [25]) | EN/RR/PLS<br>(This Work) | Our ANN<br>(This Work) | Train Mean<br>(From [7] and [18]) |
| --- | --- | --- | --- | --- | --- |
| Mean PRE ($\pm\sigma$) | 16.25±18.76 | 15.51±10.40 | 24.82±20.39 | 2.35±2.75 | 36.47±18.57 |
| Median PRE | 13.73 | 13.83 | 21.66 | 1.39 | 40.75 |
| Times Best Model | 1 | 4 | 1 | 28 | 0 |

The median and mean prediction errors for the linear models are higher than for any of the ANN models. On the other hand, the linear models have the lowest prediction errors for one of the glycans, which suggests that any nonlinearity in the true relationship for that glycan is low enough that the bias of assuming linearity is small relative to the increase in variance associated with having more degrees of freedom in the training of the ANN models [19].

The PRE for each model for each glycan is reported in Table 2. All of the models have low prediction errors for some glycans (e.g., Fc-DAO-GnGnF), but other glycans tend to have higher prediction errors (e.g., Asn38-NaGnGnGnF). These prediction errors are not significantly correlated with the Train Mean prediction error (all *R*^2^ *<* 0.10, except for the EN/RR/PLS models, which have *R*^2^ = 0.51). Our ANNs are the best model 28/32 times and produce lower prediction errors than the training mean for all glycans. In contrast, the predictions for ANN-2 and ANN-3 are worse than the training mean for some glycans.

**Table 2:** Test set percent relative errors (PREs) on the main glycans for different N-glycosylation models. Models “ANN-2” and “ANN-3” came from Ref. [25]. Models “EN/RR/PLS” and “Our ANN” come from this work. For each glycan, the “EN/RR/PLS” column reports the PRE for the model type with the best cross-validation score. “Train mean” is the PRE when using the mean of the training data (Refs. [7] and [18]). For each glycan, the lowest PRE(s) among the models is in bold. “N/A” marks predictions that are not reported in Ref. [25].

| Glycan | ANN-2<br>(From [25]) | ANN-3<br>(From [25]) | EN/RR/PLS<br>(This Work) | Our ANN<br>(This Work) | Train Mean<br>(From [7] and [18]) |
| --- | --- | --- | --- | --- | --- |
| Asn24-GnGnF | 19.95 | 33.25 | 35.87 | **4.75** | 55.40 |
| Asn24-GnGnGnF | 1.02 | 9.48 | 16.91 | **0.44** | 44.39 |
| Asn24-GnGnGnGnF | 12.88 | **1.64** | 26.93 | 5.63 | 47.89 |
| Asn24-MGnF | 19.51 | 25.37 | 35.17 | **5.88** | 47.80 |
| Asn38-GnGnF | 3.00 | 18.39 | 29.64 | **0.56** | 43.43 |
| Asn38-GnGnGnF | 7.11 | **1.74** | 33.56 | 3.63 | 45.54 |
| Asn38-GnGnGnGnF | 19.36 | 13.10 | 52.87 | **1.79** | 60.82 |
| Asn38-NaGnGnGnF | 51.70 | **5.68** | 89.20 | 13.56 | 45.45 |
| Asn83-GnGnF | 20.11 | 25.00 | 42.61 | **2.44** | 59.92 |
| Asn83-GnGnGnF | 3.41 | 8.92 | 40.48 | **0.90** | 57.84 |
| Asn83-GnGnGnGnF | 5.59 | 14.55 | 57.73 | **2.30** | 65.07 |
| Asn83-NaGnGnGnF | 100.00 | 20.00 | 52.22 | **0.00** | 22.78 |
| Asn110-Man5 | 27.07 | N/A | 12.01 | **0.00** | 10.26 |
| Asn110-Man6 | 5.43 | N/A | 3.94 | **0.67** | 4.02 |
| Asn110-Man7 | 27.78 | N/A | 8.54 | **0.51** | 16.54 |
| Asn168-GnGnF | 24.53 | 27.79 | 36.41 | **0.00** | 56.19 |
| Asn168-GnGnGnF | 17.12 | 19.62 | 33.10 | **5.00** | 57.61 |
| Asn168-GnGnGnGnF | 23.15 | 28.64 | 42.36 | **3.67** | 63.86 |
| Asn168-MGnF | 10.81 | 6.91 | 26.17 | **1.18** | 39.56 |
| Asn538-GnGn | 21.80 | 22.56 | 25.94 | **1.13** | 45.80 |
| Asn538-GnGnF | 6.80 | 5.65 | 10.99 | **3.81** | 36.82 |
| Asn538-MGn | **1.59** | **1.59** | 12.64 | **1.59** | 27.65 |
| Asn538-MGnF | 3.88 | 4.71 | 1.11 | **0.00** | 19.04 |
| Asn745-GnGn | 15.67 | 10.26 | 17.38 | **0.57** | 41.93 |
| Asn745-GnGnF | 6.32 | 7.48 | 10.02 | **1.62** | 36.12 |
| Asn745-MGn | 17.21 | 32.56 | 11.35 | **5.56** | 21.28 |
| Asn745-MGnF | 19.20 | 27.23 | **2.19** | 2.67 | 15.29 |
| Fc-DAO-GnGnF | 2.73 | N/A | 3.03 | **1.00** | 19.12 |
| Fc-EPO-GnGnF | 1.36 | N/A | 4.46 | **0.11** | 21.71 |
| modelNSD-GnGnF | 7.85 | N/A | 9.51 | **0.74** | 14.63 |
| modelNSD-AGnF | 1.36 | N/A | 2.97 | **0.95** | 6.40 |
| modelNSD-AAF | 14.57 | N/A | 7.07 | **2.60** | 16.98 |

The distribution of prediction errors for our ANNs are concentrated near low values (Fig. 2). At the other extreme, the distribution of prediction errors for ANN-2 has a wide spread with very high prediction error (*≥* 50%) for two glycans. The linear models also have a long tail, although not to the same degree. ANN-3 has a lower range of prediction errors, but it is still significantly higher than that of our ANNs (Fig. 2).

**Figure 2:**
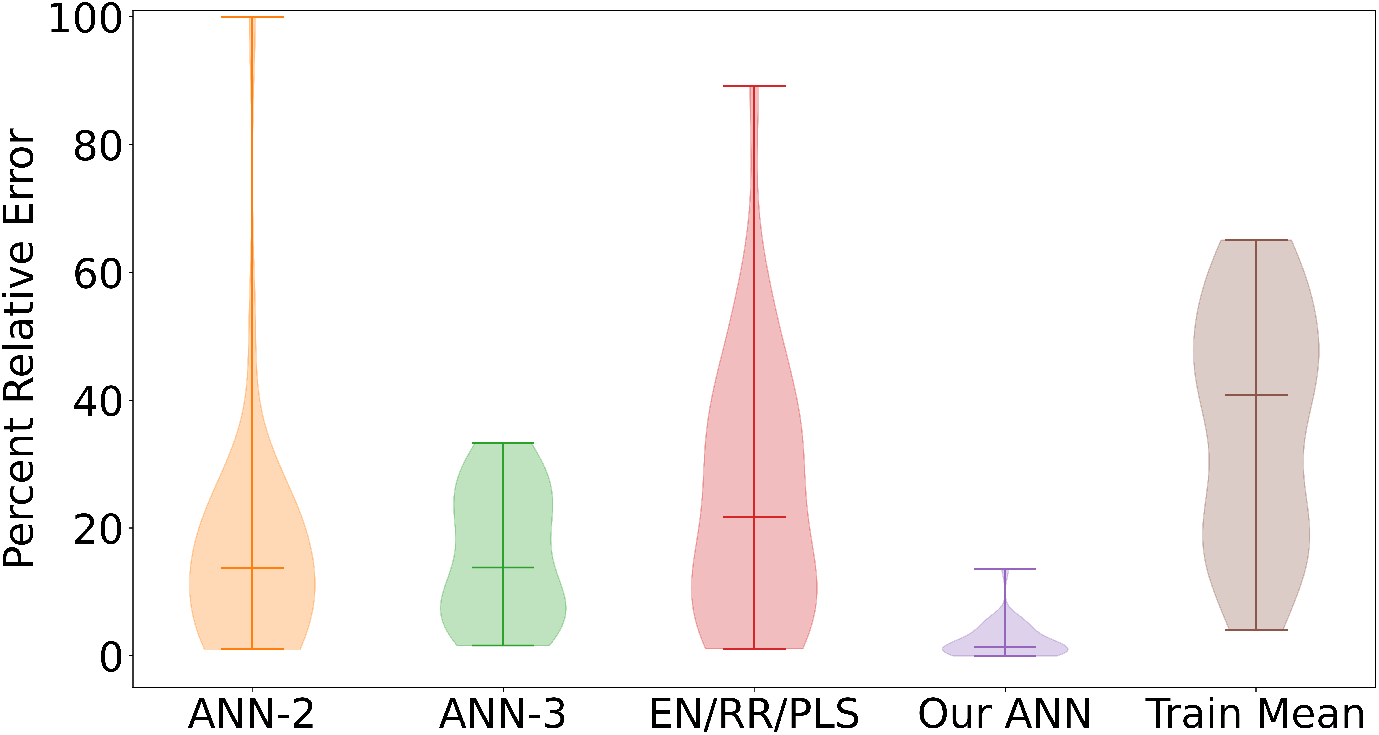
Violin plots showing the distribution of test set PREs for different models. Model labels are as in Fig. 1. The horizontal bars within the distributions are median PREs.

### 3.2 Nested validation highlights the robustness of our models

This work uses the same test set as Ref. [25] to allow a fair comparison between models. To ensure the models constructed in this study are not biased by that choice of test set, nested validation was performed. In each round, a group within the complete dataset is selected as the test set, and the rest is used for training and cross-validation. These rounds repeat until all groups have been used as the test set. In this work, five rounds of nested validation are performed, each following the procedure in Section 2.1.

The difference is negligible between the PREs of the models selected with and without nested validation (Fig. 3). The Jensen-Shannon distance between the EN/RR/PLS distribution is 0.127 and that between the ANN distributions is 0.478. As such, the selection of test set did not introduce any significant bias to the models, and the models are robust to changes in the split of data into training and testing.

**Figure 3:**
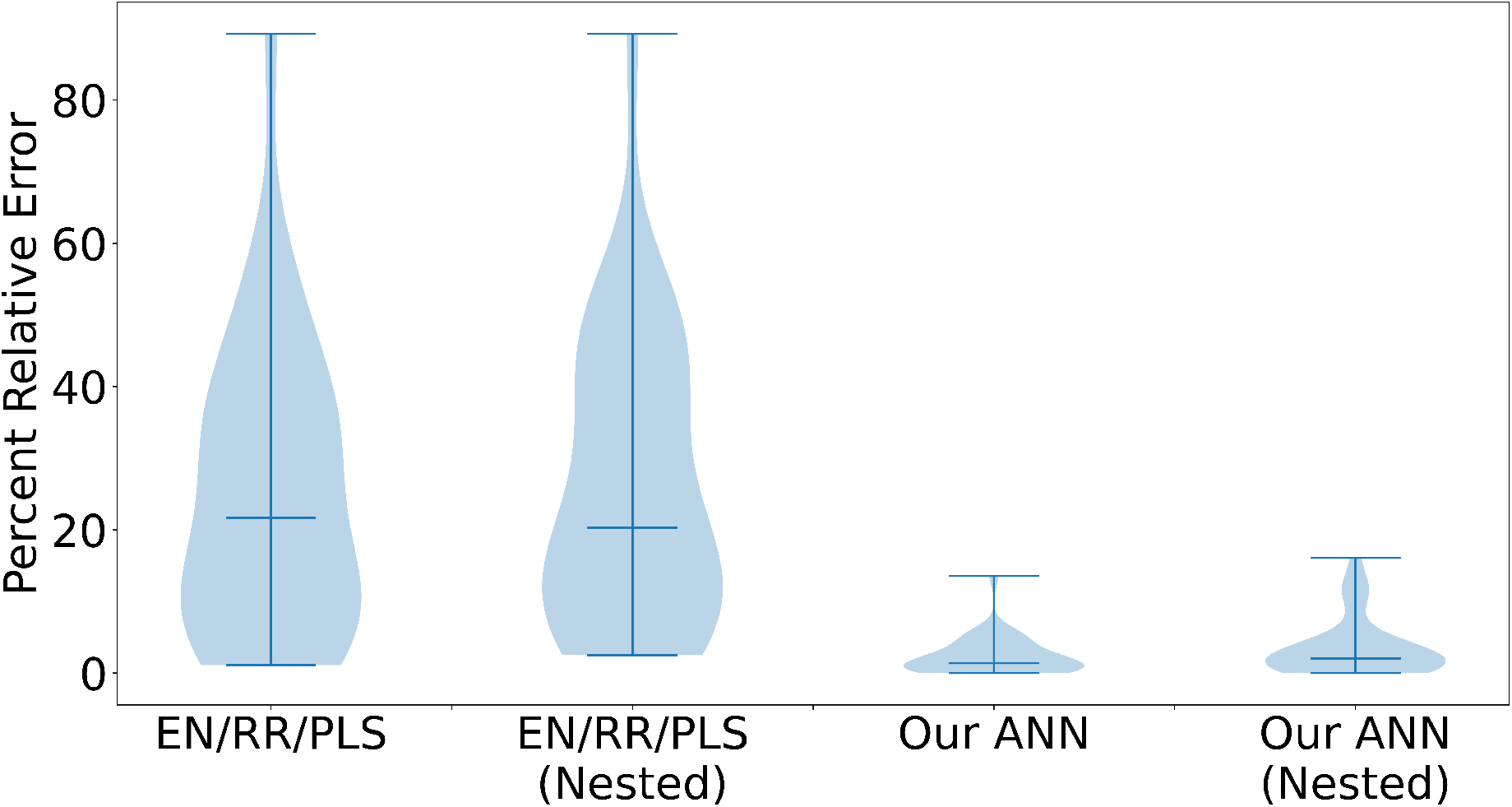
Violin plots showing the distribution of test set PREs before and after nested validation.

## 4 Discussion

This study constructs models from literature data on antibody and fusion protein glycan distribution based on normalized CHO cell B4GALT1–B4GALT4 levels. Two types of models are trained to predict the glycan distribution based on user-input B4GALT levels. Linear models (EN/RR/PLS) provided reasonable predictions of glycan distribution despite their simplicity. Artificial neural network models are constructed that have significantly improved prediction performance compared to published models. The median PRE of the new ANN models is 1.39%, with the error distribution concentrated near low values. This median PRE is 10-fold lower than for the other models. A nested validation study shows that the prediction performance of the linear and ANN models was insensitive to the split of the data into training/validation and test sets.

The models trained in this work are compared to the ANN models of Ref. [25], which did not properly separate the experimental data into training, validation, and test sets. This issue effectively ensured Ref. [25]‘s models are overfit, resulting in overly optimistic estimates of the prediction errors. Ref. [25] manually set the ANN structures, instead of using a pre-published package such as PyTorch as in this study, and fixed the choice of some hyperparameters, namely, the learning rate and activation function. The models in Ref. [25] also were not trained using learning rate scheduling. These choices limit the potential prediction performance of the models in Ref. [25], and explain why the PREs of their models are high despite being overfitted to the test data.

This software used in this work is publicly available,^5^ allowing other researchers to reproduce this work and reuse or improve the code in future studies. The software provides a simple way to install and run the models to predict the glycan distribution of antibodies and fusion proteins made in CHO cells given normalized B4GALT1–B4GALT4 levels.

## Acknowledgements

This study was financially supported by a Project Award Agreement from the National Institute for Innovation in Manufacturing Biopharmaceuticals (NIIMBL), U.S., with financial assistance from awards 70NANB17H002 and 70NANB20H037 from the U.S. Department of Commerce, National Institute of Standards and Technology. P.S. was partially supported by a MathWorks Engineering Fellowship.

## Appendix

### Reproducing the models and plots

The nested and regular cross-validation results can be recreated by downloading the datasets and running the `SPA_glycosylation_model.py` file without any flags (`python SPA_glycosylation_model.py`) and with the --nested flag (`python SPA_glycosylation_model.py --nested`). The ANN results can be recreated by downloading the `ANN_train.ipynb` file, changing the first cell as needed, and running the notebook.

The plots can be recreated by downloading the `result_csv_files` and `result_csv_files_nested` folders, then running the `make_results_plots.py` file. Most plots will be generated in the first folder, and the nested validation PRE distribution will be generated in the nested results folder.

### Using the models to predict glycan distributions

The Conda environment defining the specific packages and version numbers used in this work is available as `ANN_environment.yaml` on our GitHub (github.com/PedroSeber/CHO_N-glycosylation_prediction). To use our trained models, run the `ANN_predict.py` file as `python ANN_predict.py <location>` `<B4GALT levels>`. For example, use the command `python ANN_predict.py Asn_24 1 1 1 1` to predict the wild-type glycan distribution at Asn 24, or `python ANN_predict.py Asn_83 0.001 0.0041.03 1.1` to predict the glycan distribution at Asn 83 of a double-knockout mutant.

## Footnotes

1 All data are available as .csv files on our GitHub (github.com/PedroSeber/CHO_N-glycosylation_prediction).

2 Group labels are available in the group_names.txt and group_names_NN.txt files on the GitHub repository.

3 The software version used in this article is available at github.com/PedroSeber/SmartProcessAnalytics.

4 As detailed in the ANN_train.ipynb file in the GitHub repository.

5 https://github.com/PedroSeber/CHO_N-glycosylation_prediction.

